# Microbial partners drive legume trait plasticity and tripartite interaction outcomes under combined stress environments

**DOI:** 10.1101/2025.10.02.678283

**Authors:** Brendan A. Randall, Kimberly J. Komatsu, John D. Parker, Kelsey McGurrin, Sarah J. M. Alley, Chase T. Hearn, Karin T. Burghardt

**Author notes:** Corresponding author: Brendan A. Randall.

## Abstract

Plants often form partnerships with symbiotic microbes that improve plant performance. Increasing environmental stress is changing interaction dynamics between plants and their below-ground symbiotic partners, which can have cascading effects on above-ground trophic interactions. However, little work has examined how microbes protect plants against multiple stressors acting in concert, or the extent to which microbial genetic variation shapes context-dependent outcomes of tripartite interactions between microbes, plants, and herbivores. We experimentally manipulated the genetics of symbiotic microbial partners by individually inoculating a single legume species (*Glycine max*) with twenty-four genetically distinct strains of rhizobial bacteria. We quantified insect herbivore feeding and growth rates, as well as plant ecophysiological and performance traits under different combinations of drought and herbivore exposure. Rhizobial strains differentially influenced the effect of drought on legume ecophysiological traits. Certain strains increased photosynthesis and enhanced photoprotection under drought. Moreover, legumes under drought and herbivore exposure also exhibited increased photoprotection. Some strains were observed to limit herbivore growth rates under drought more effectively than others. A multivariate trait analysis revealed that strains differentially influenced legume trait syndromes and plasticity in response to stress in certain environments. Rhizobial strain variation was a key driver of tripartite interactions through changes in plant trait expression and plasticity under simultaneous abiotic and biotic stress, which had cascading effects on insect herbivore growth. A community approach incorporating multiple beneficial microbial strains may be effective for managing legume-rhizobia symbioses and introducing functional trait diversity and resistance traits into ecosystems.

## Introduction

Understanding how biological communities are changing due to increasingly variable and extreme environments requires a system-level approach that incorporates multitrophic interactions. Although pairwise species interactions underly important biological processes such as food web dynamics, primary productivity, and nutrient cycling (Paine, 1966; Schmitz et al., 2000; Vitousek et al., 2013), they occur within a broader context of abiotic factors and other biological interactions (Stanton, 2003; Heath & Lau, 2011; Chamberlain et al., 2014). For example, below-ground symbiotic microbial partners can alter above-ground plant plasticity to environmental stress (Denison & Kiers, 2004; Goh et al., 2013; Bolin, 2025). Increasing variability in abiotic and biotic environments has also been shown to shift the frequency and intensity of insect herbivory due to changes in plant host quality and defense, with consequences for arthropod community structure (Grinnan et al., 2013; Wetzel et al., 2016; Bertolet et al., 2024; Cope & Wetzel, 2025). However, the importance of below-ground microbial symbionts in shaping the sensitivity of above-ground plant and insect responses is rarely examined in the context of combined abiotic and biotic stressors (Chamberlain et al., 2014; Hoeksema & Bruna, 2015).

The rhizobia-legume system is a common model for understanding below-ground interactions mediate above-ground interactions. Rhizobia are a group of specialist nitrogen-fixing bacteria (Family: Rhizobiaceae) that colonize roots and form symbiotic, often mutualistic interactions with leguminous plant hosts (Family: Leguminosae; Fabaceae). Plants benefit by receiving fixed, usable nitrogen (N) from rhizobia in exchange for carbon (C) and protection within root nodules that facilitate rhizobial reproduction (Sprent, 1972; Somasegaran & Hoben, 1994; Vance, 2001; Udvaldi & Poole, 2013; Poole et al., 2018). As legumes are key crops around the world, these symbioses are critical for safeguarding the global food supply and for legume crop production in stressful environments (Siambele et al., 2021).

The genetic pool of rhizobial bacteria in soils is exceptionally diverse, comprised of many distinct genotypes, or strains (Brockwell et al., 1995; Koppell & Parker, 2012; Taylor et al., 2020). Rhizobial strains vary in their tolerance to environmental stressors such as drought (Zahran, 1999; Heath & Tiffin, 2007; Ngumbi & Kloepper, 2016), and can also positively or negatively influence insect herbivore performance (Bronstein & Barbosa, 2002; Heath & Lau, 2011; Ballhorn et al., 2013; Thompson & Lamp, 2021).

Certain rhizobial partners provide legumes with consistent, high levels of N that mitigate stress in variable environments (Sprent & Sprent, 1990; Yuan et al., 2020), which can improve leaf tissue quality and availability that promotes herbivory (Salvagiotti et al., 2008; Couture et al., 2010; Dean et al., 2014). Conversely, legumes can benefit by incorporating this N into chemical defenses (Nahrstedt, 1985; Koricheva et al., 2004; Bhattacharyya & Jha, 2012) that inhibit herbivory (Ballhorn et al., 2007; Kempel et al., 2011; Thamer et al., 2011). However, these tradeoffs vary by strain and influence symbiotic effectiveness and herbivore growth rates, which may shift under stress (Denison & Kiers, 2004; Heath & Tiffin, 2007; Heath & Lau, 2011). Therefore, performance of plant-rhizobial partnerships may depend on individual strain tolerance to abiotic and biotic conditions (Brockwell et al., 1995; Prudent et al., 2020).

Trait-based experimental approaches may reveal how the genetics of individual rhizobial strains modulate plant-insect interaction outcomes. Plant functional traits impact fitness by affecting plant growth, reproduction, and survival (Cornelissen et al., 2003; Violle et al., 2007; Liu et al., 2021) and, therefore, serve as parameters to examine symbiotic benefits. For example, rhizobial-plant partners engaged in optimal trading of N lead to increased photosynthetic rates (Peng et al., 2002; Govindasamy et al., 2017) and biomass production (Jensen & Hauggard-Nielsen, 2003). Nodule abundance can indicate the effectiveness of partner colonization (Lindström & Mousavi, 2020), which correlates with improved N fixation and overall plant fitness (Pereira et al., 1993; Batstone et al., 2022; Zhong et al., 2024). However, it remains unclear whether strain-specific differences on plant functional traits can indirectly drive variation in plant resilience to stress.

Here, we used a combination of trait-based approaches and herbivory assays in a rhizobial strain manipulation greenhouse experiment to examine genetics-by-environment interactions, specifically whether environmental stress modifies the above-ground plant-insect interaction outcomes with specific below-ground microbial partners (Fig. 1). First, we ask whether there are strain-specific differences in caterpillar herbivore performance on host plants under drought, testing for an interaction of rhizobial genetics-by-environment (G_Strain_×E_Drought_). We predict that rhizobial strains will differentially influence caterpillar growth rates and herbivory during drought. Therefore, we predict that rhizobial genetics will have a cascading effect on herbivore growth, which will be modified by drought. Next, we ask if there are strain-specific differences in plant traits under simultaneous drought and herbivore stress. We predict that plant trait responses to drought and herbivory differs in response to association with different rhizobial strains (G_Strain_×E_Drought_×E_Herbivory_). Finally, we ask whether rhizobial strain identity affects plant performance and resource allocation patterns, and to what extent these factors respond to abiotic and biotic stressors. We predict that plants in well-watered, herbivore-free environments (E_Drought_×E_Herbivory_) will have greater nodule counts and biomass with reduced root:shoot ratios due to shifting resource allocation towards leaf tissues in response to reduced environmental stress. This research highlights the dynamic, context-dependent nature of provider mutualisms, and supplies a framework for future research investigating how genetic and environmental variation shapes ecological interactions.

**Figure 1.**
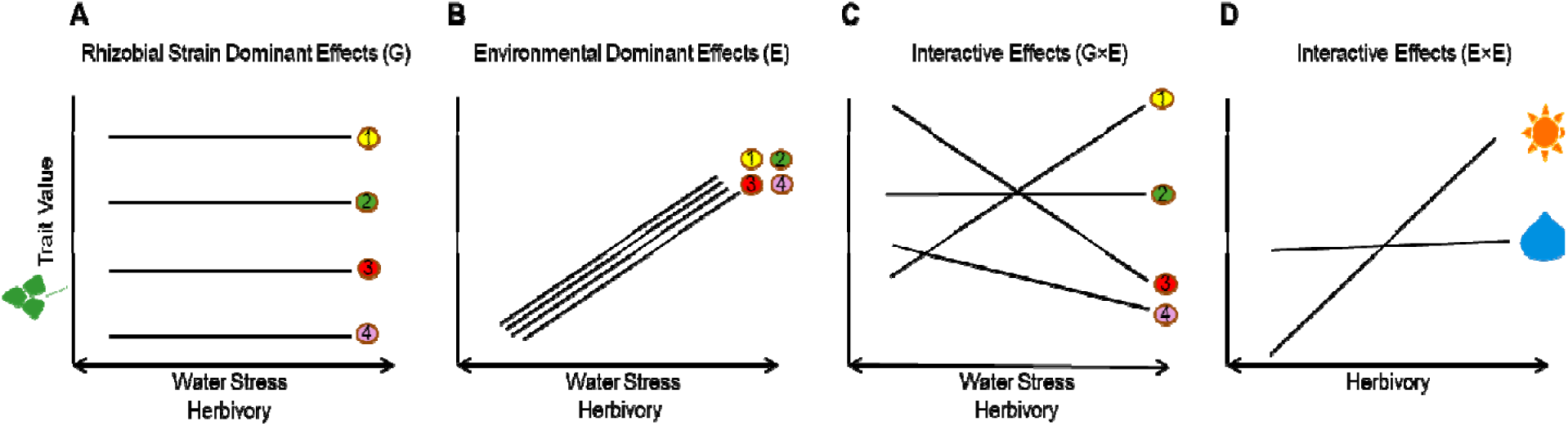
A conceptual diagram for understanding rhizobial (G) and environmental (E) dominant effects, as well as their interactions (G×E and E×E), on plant trait expression and plasticity in combined abiotic and biotic stress environments: water stress (raindrop-well-watered; sun-drought) and herbivory. (A) Genetically distinct rhizobial strains (colored dots; 1-4) drive unique but consistent plant trait expression and low trait plasticity across all environments. (B) Environmental dominant effects (E) drive plant trait expression and plasticity. For example, plant trait values increase in both drought and herbivore environments, but do so almost equally between all strains. (C) Genetic variation exists for resistance trait plasticity, in which the effect of watering or herbivory on plant trait expression is dependent on the specific strain a plant was inoculated with (G×E). Certain strains increase plant trait expression under stress, while other strains decrease trait expression. (D) The effect of one environment changes depending on another environment. For example, the effect of herbivory on plant trait values is greater in droughted plants than watered plants.

## Materials and Methods

### Rhizobial trapping experiment and strain selection

Soybean (*Glycine max*) was chosen as our model legume plant species because it is among the most common legume crops grown globally, but its production is increasingly threatened by drought and recurring pest outbreaks (Rosenweig et al., 2001; Cornelissen et al., 2011; IPCC 2014). In June 2020, we planted an untreated soybean variety (cv. ‘Dekalb Asgrow AG38X8’) in three 5.5 m x 3.5 m experimental plots at the University of Maryland Central Maryland Research and Education Center (CMREC), in Clarksville, MD (39.2552° N, 76.9292° W). Three weeks after planting, four soil subsamples were collected from each plot and refrigerated until spring 2021. These samples were then used as soil slurry inoculants for a rhizobial trapping experiment completed in spring 2021. For this trapping experiment, we grew inoculated soybeans of the same cultivar in a field trial (*N* = 234 plants) under ambient or droughted conditions. Plants in the ambient and drought treatments were watered twice daily in 15-minute intervals during the first two weeks of growth, after which watering was reduced to ten minutes every 12 hours in the ambient treatment and five minutes in the droughted treatment. Plants were harvested eight weeks after planting. Rhizobial strains were trapped by first collecting ten surface-sterilized nodules from each plant. Cells were then isolated and cultured. To identify rhizobial species from all resultant cultures, we sequenced the *nifH* gene region, the most widely used biomarker gene for nitrogen-fixing bacteria (Raymond et al., 2004; Gaby & Buckley, 2014). Isolated strains were identified into 123 unique rhizobial genotypes. From these, we selected 24 unique strains (*N* = 24) based on their genetic distance, sample abundance, and preference for either the well-watered or droughted environment (Fig. S1). The selected rhizobial strains were classified within the *Bradyrhizobium diazoefficiens* (*N* = 21) and *Bradyrhizobium elkanii* (*N* = 3) species complexes.

### Experimental design

To understand the role of combined stressors on the ability of rhizobial strains to mediate plant-insect interaction outcomes, we completed a greenhouse experiment in summer 2022 at the University of Maryland Research Greenhouse Complex (38.9976° N, 76.9378° W). Soybean seeds (cv. ‘Dekalb Asgrow AG38X8’) were surface sterilized with bleach and germinated in the lab for three days. One sprouted seed was then planted into each of 624 6 cm x 25 cm ‘Cone-tainer’ pots (Stuewe & Sons, Inc., Tangent, Oregon) filled with sterile sand.

Plants were arranged in a systematic random block design consisting of six replicate blocks, with each replicate being divided into: 24 strain treatments, two watering treatments (well-watered and drought), and two herbivory treatments using a leaf-chewing caterpillar whose preferred host is soybean (Martin et al., 1976), *Chrysodeixis includens* (Lepidoptera: Noctuidae) (+ herbivory: caterpillar present, - herbivory: caterpillar absent). Each strain, drought, and herbivory combination contained six plants (24 strain treatments X 2 watering treatments X 2 herbivore treatments X 6 replicates - 576 experimental plants). Plants were alternately assigned a well-watered or drought treatment to account for inter-block variation in greenhouse climate conditions throughout the day (*N* = 312 plants/watering treatment), with each replicate block consisting of eight trays, each containing 12 plants. No strain, drought, and herbivory combinations were replicated in the same block. Additionally, one plant not inoculated with rhizobia was positioned in the middle of each tray to serve as a negative control (*N* = 48 control plants) to monitor cross-contamination between plants within each tray. Nodulation was not observed among these control plants, and therefore cross-contamination among inoculated plants was considered inconsequential. Within each tray, two plants assigned to the same strain and watering treatment were paired in one of the two herbivory treatments (*N* = 288 plants/herbivory treatment). The spatial order of rhizobial strain treatments was randomized within each replicate block.

Liquid rhizobial culture of each of the 24 strains selected for this experiment were grown in Yeast Mannitol Broth to a density of roughly 1 million cells per mL, estimated by optical density of 0.26 at 600 nm. Three weeks post-emergence, treatment plants were inoculated with 1 mL of liquid rhizobial culture from one of the twenty-four rhizobial strains administered via repeating pipette to the base of the plant. During the first three weeks post-emergence, all plants were watered four times daily for 60 minutes total. At one-week post-inoculation, plants in the ambient ‘well-watered’ benches were watered four times daily for 60 minutes total, while plants in the ‘drought’ treatments were watered twice daily for 30 minutes total. This regimen was established based on qualitative observations of wilting of drought-induced stress on plants.

### Caterpillar feeding trial

Six weeks post-emergence, we completed an herbivore feeding trial with two goals. The first goal was to test whether rhizobial strain identity influenced plant resistance to *C. includens* larvae. Second, the feeding trial also served as a biotic stress imposed on the plants to understand whether herbivory induces later changes in plant trait expression or resource allocation. We purchased third-instar *C. includens* larvae from a commercial insectary (Benzon Research, Carlisle, PA), which were reared on an artificial diet. Upon arrival, individual caterpillars were food-deprived for 12 hours, weighed, and placed on a single trifoliate leaf of a plant assigned to the ‘+ herbivory’ treatment, contained within a mesh bag. Caterpillars fed on the plants for 36 hours, then were removed, food-deprived for 12 hours, and weighed again to calculate relative growth rates (weight_final_ - weight_initial_) ÷ (weight_initial_). After caterpillar feeding, we quantified total herbivory using LeafByte, a mobile image analysis application that calculates the proportion and total amount of leaf area removed by herbivores (Getman-Pickering et al., 2020). We removed entire trifoliate leaves damaged by caterpillars (about 25% of total plant biomass) after the feeding trial to standardize damage across all ‘+ herbivory’ plants.

### Plant traits

Plant trait data was collected at eight weeks post-emergence to assess variation in response to rhizobial strain identity, abiotic (drought), and biotic (herbivory) stress.

First, we harvested leaf disk tissue samples (area = 19.63 mm^2^) from the upper-most fully expanded trifoliate leaf. Disks were immediately weighed, dried, and reweighed to calculate specific leaf area (standard leaf area (mm^2^) ÷ dry weight (mg)) and percent relative water content ((fresh weight (mg) – dry weight (mg)) ÷ fresh weight (mg) X 100) (Vile et al., 2005). At nine weeks post-emergence, we used a MultiSpeq PhotosynQ (Kuhlgert et al., 2016) to simultaneously obtain multiple focal photosynthetic and photoprotection indicator traits: Φ*_II_* (estimate of maximum plant photosynthetic yield; Kramer et al., 2004; Kuhlgert et al., 2016), Φ*_NO_* (estimate of excess light energy dissipated by the plant through non-regulatory heat loss; Kramer et al., 2004; Kuhlgert et al., 2016), NPQt (estimate of plant capacity to dissipate excess light energy as heat; Kuhlgert et al., 2016; Tietz et al., 2017), relative chlorophyll content, and leaf temperature differential. We also measured plant shoot height and the number of fully formed trifoliate leaves for each plant. Ten weeks post-emergence, plants exhibited visible signs of senescence and were destructively harvested. Roots were washed and nodules from each plant were counted. The number of developed pods were counted, and all plant parts (shoots, roots, and pods) were dried and weighed for biomass and resource allocation metrics.

### Statistical analysis

All statistical analyses were conducted in R version 4.1.0 (R Core Team, 2021). To examine the effect of rhizobial strain and drought on herbivores, we performed linear mixed effect models using the *lmer* command in the *lme4* package (Bates et al., 2015). Response variables were caterpillar relative growth rates and percentage leaf area consumed; rhizobial strain identity, drought, and their interaction (G×E) were fixed effects. To determine the effects of rhizobial strain, drought, and herbivory on plant functional trait responses, we used linear mixed-effect models for the following response variables: specific leaf area, percent leaf water content, Φ*_II_*, Φ*_NO_,* NPQt, relative chlorophyll, leaf temperature differential, shoot biomass, root biomass, root:shoot ratio, plant height, and nodule count (Table S1 and S2). We used rhizobial strain, drought, and the presence of herbivores as fixed effects, including all possible two-way interactions. A random intercept for replicate was included to account for our systematic randomized block design. Hypothesis testing was conducted using Type III sums of squares with degrees of freedom estimated using the Satterthwaite approximation in the *lmertest* package (Kuznetsova et al., 2017). For the description and interpretation of results, we considered strong evidence of effects where *p* < 0.05, and weak evidence where 0.05 < *p* < 0.10 (Muff et al., 2022) (Table S2).

Next, we used a constrained ordination approach, redundancy analysis (RDA, Oksanen et al., 2022) using Euclidean distances to examine changes in whole-plant trait expression in response to our experimental treatments. We used the *rda* command in the *vegan* package to build a model on twelve measured functional traits (Table S3), which were centered and standardized across traits using the *decostand* function (Oksanen et al., 2022). We included rhizobial strain, drought, and the presence of herbivores as fixed effect predictor variables, and all possible two-way interactions. A conditioning term removed variation associated with the systematic randomized block design of the experiment. A permutational pseudo-F-test using the *anova* command assessed model fit. Hypothesis testing for individual fixed effects was only performed if the overall model was a good fit for the data. To assess potential differences in dispersion between centroids in each watering treatment, we used the *betadisper* function.

## Results

### Does herbivore performance vary by rhizobial strain and drought?

Herbivores grew at different rates depending on the rhizobial strain and watering treatment (Fig. 2; F_23,231_ = 1.55, *p* = 0.06). Rather than a uniform response, certain strains were associated with higher caterpillar growth in well-watered environments, but lower under drought (Fig. 2; negative slopes), and vice versa (Fig. 2; positive slopes).

**Figure 2.**
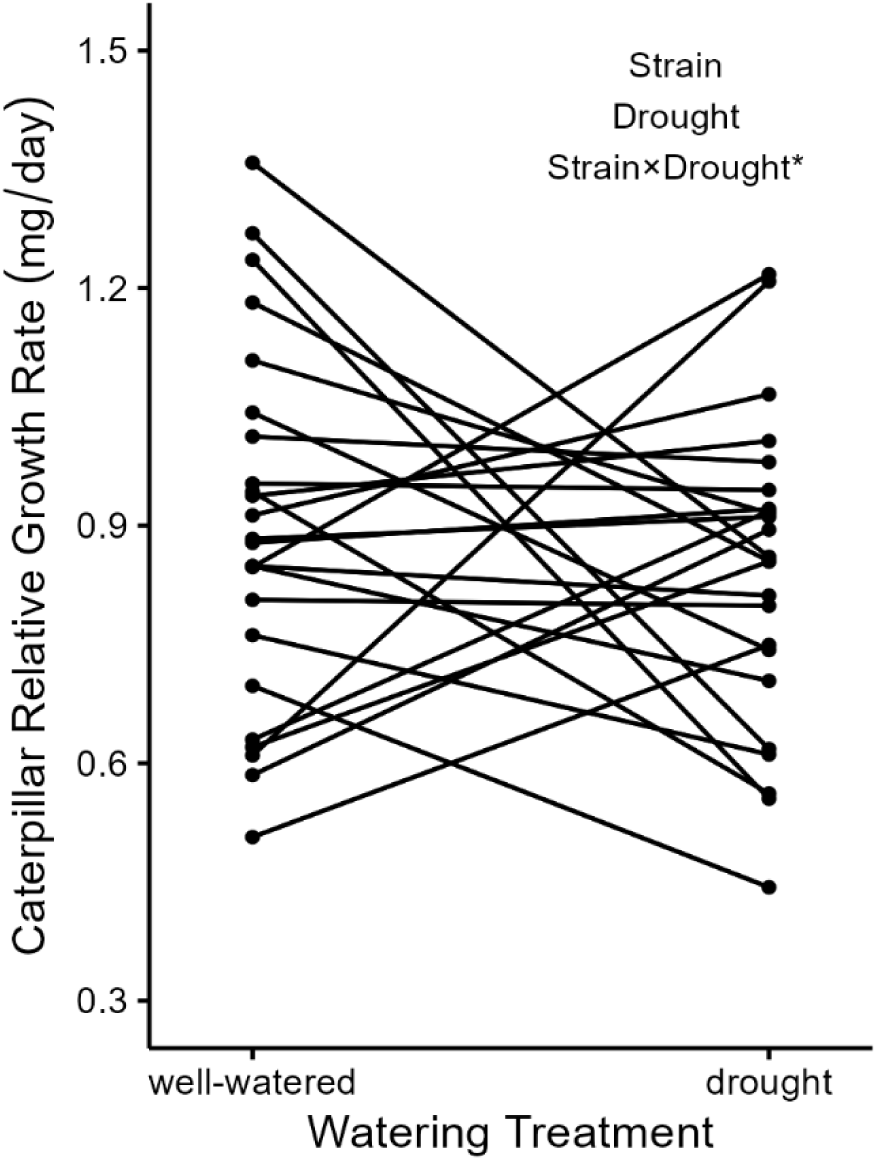
Rhizobial strain influences drought-mediated herbivore growth rates. Reaction norms represent the mean ± standard error (SE) effect of watering (well-watered or drought) on caterpillar relative growth rate. Each line represents one of 24 rhizobial strain genotypes in which 12 plants were inoculated in either well-watered or droughted conditions. * = 0.05 < p < 0.10.

However, we found no effects of strain, watering, or interactive genetic and environmental effects on caterpillar herbivory (% leaf area consumed; Table S2).

### How do plant functional traits respond to rhizobial strain, drought, and herbivory?

Genetic and environmental effects often interactively determined plant functional trait expression. The effect of drought on maximum photosynthetic yields (Φ*_II_*) was dependent on rhizobial strain (Fig. 3A; F_23,482_ = 2.25, *p* < 0.001). Certain strains enhanced or maintained Φ*_II_* under drought (Fig. 3A; positive or parallel slopes), while others decreased Φ*_II_* (Fig. 3A; negative slopes). Additionally, ‘+ herbivory’ plants had 3% higher Φ*_II_*than ‘-herbivory’ plants (Fig. 3B; F_1,482_ = 4.49, *p* = 0.026). The effect of drought on non-photochemical quenching (NPQt) was also dependent on rhizobial strain (Fig. 3C; F_23,482_ = 2.20, *p* < 0.001). NPQt was 20% lower in ‘+ herbivory’ plants compared to ‘-herbivory’ plants (Fig. 3D; F_1,482_ = 7.47, *p* = 0.007). We also observed a significant G_Strain_×E_Drought_ interaction on plant non-regulatory heat loss (Φ*_NO_*), with certain strains increasing expression of Φ*_NO_* under drought, with others decreasing Φ*_NO_* (Fig. 3E; _F23,482_ = 2.35, *p* < 0.001). Among droughted plants, ‘+ herbivory’ plants were less effective at regulating heat loss, with 12% greater Φ*_NO_*compared to ‘-herbivory’ plants (Fig. 3F; F_1,482_ = 4.41, *p* = 0.042).

**Figure 3.**
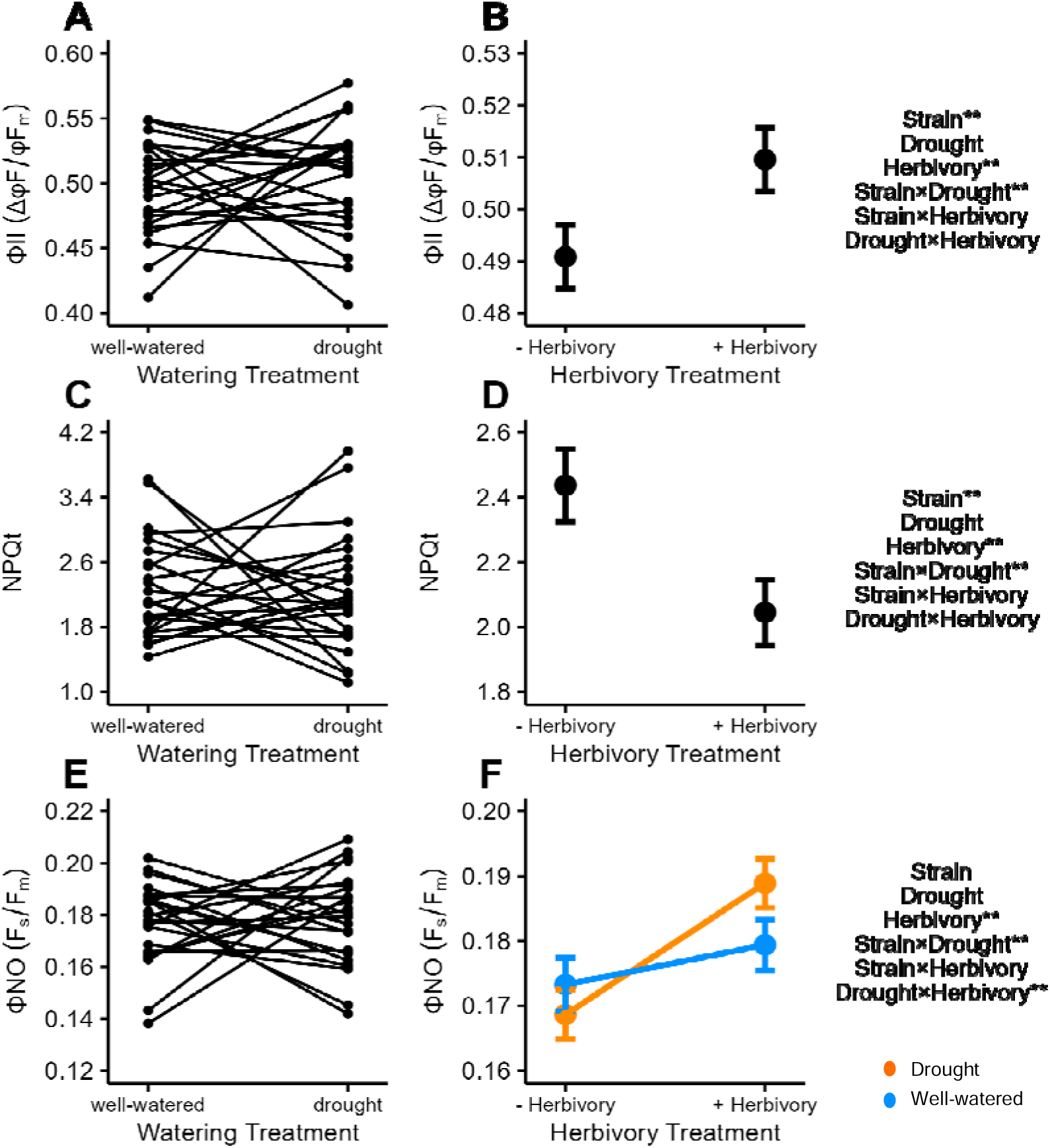
Effects of rhizobial strain identity, watering, herbivory, and their interactions on plant photosynthetic and photoprotection traits (A-F). (A) Mean Φ_II_ ± standard error (SE) reaction norm plot by each of the 24 rhizobial strain genotypes grouped by watering treatment (N = 12 plants/strain/watering treatment), and (B) herbivory (N = 288 plants/herbivory treatment). (C) Mean NPQt ± standard error reaction norm plot by each of the 24 rhizobial strain genotypes grouped by watering treatment (N = 12 plants/strain/watering treatment), and (D) herbivory (N = 288 plants/herbivory treatment). (E) Mean Φ_NO_ ± SE reaction norm plot by each of the 24 rhizobial strain genotypes grouped by watering treatment (N = 12 plants/strain/watering environment), and (F) by each watering treatment grouped by herbivory (N = 184 plants/watering/herbivory treatment). ** = p ≤ 0.05.

### How does whole-plant trait expression and plasticity respond to rhizobial strain, drought, and herbivory?

A redundancy analysis (RDA) assessing multivariate plant trait responses (Fig. 4A: arrows) to rhizobial strain, drought, and herbivory predictors explained 14% of total plant trait expression variation and 52% of the constrained variation (Whole model Pseudo-F_1,48_ = 1.64, *p* = 0.001). Photosynthetic and photoprotection traits separated along the first RDA axis (Φ*_II,_* Φ*_NO,_* NPQt, leaf temperature differential), accounting for 33% of plant trait variation. The second RDA axis was associated with growth and resource allocation traits (shoot biomass, root biomass, plant height, nodule count, root:shoot ratio, specific leaf area, percent leaf water content, and chlorophyll content) accounting for 19% of the variation.

**Figure 4.**
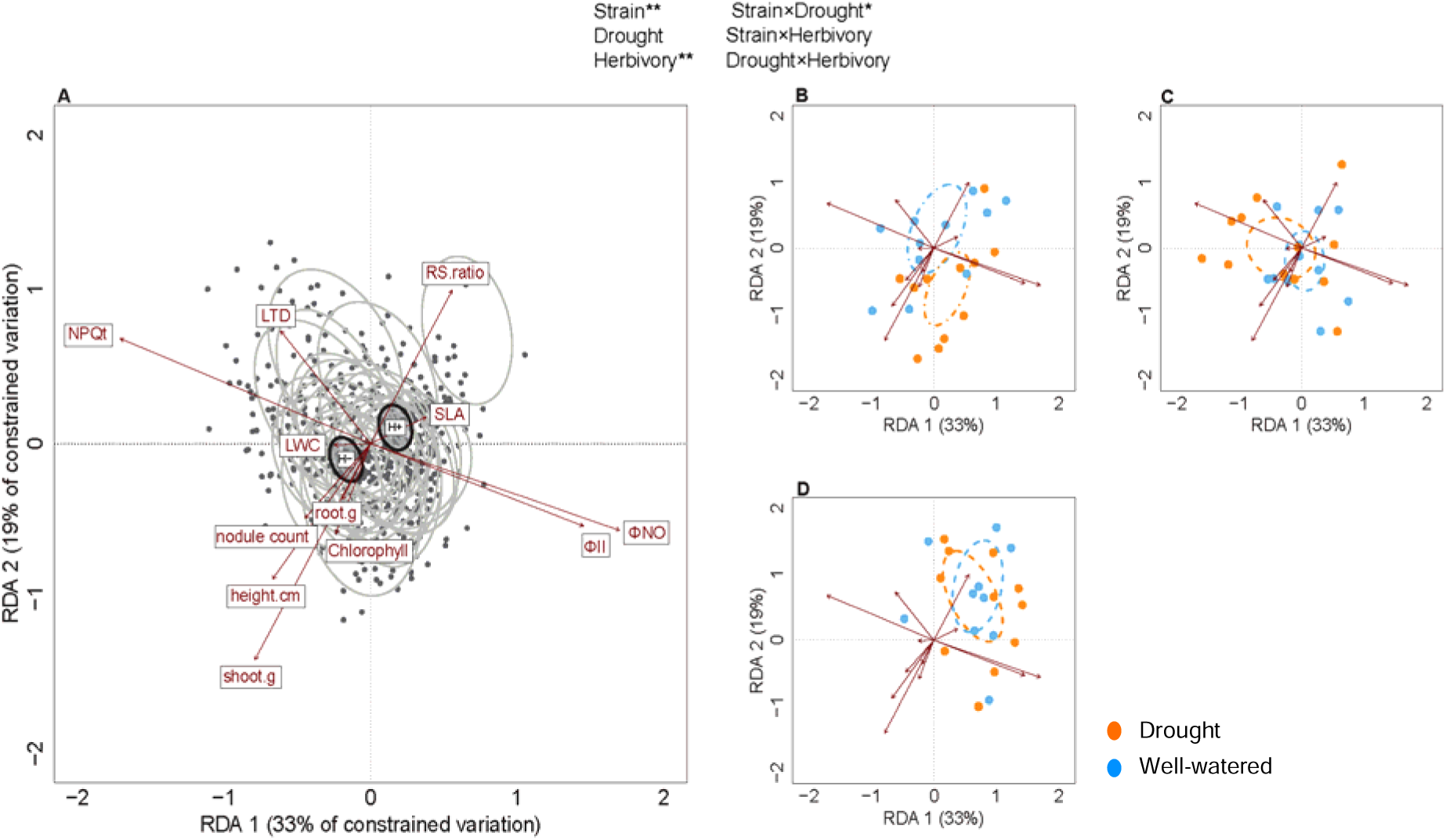
(A) Variation in average multivariate plant trait expression in response to rhizobial strain and herbivory treatments. The first two axes represent the constrained variation of an RDA plotting multivariate plant trait expression in response to each strain averaged across all environments (grey ellipses; N = 24), and herbivory treatment (black ellipses; + herbivory and - herbivory). Each dot represents an individual plant’s collection of traits. Each vector represents a response variable, a plant trait, and the association that trait has with each of the two primary axes. The origin, or centroid of each ellipse represents the weighted averages (95% CI) of every plant inoculated with one of twenty-four strains of rhizobia (N = 24 plants/strain), and each plant exposed or not exposed to herbivory (N = 288 plants/herbivory treatment). (B-D) Variation in average multivariate plant trait expression and multivariate trait dispersion in response to rhizobial strain and watering environment. The trait space is replicated from (A). Colored dots represent an individual plant’s collection of traits in a specific strain and a specific environment. The origin of ellipses represents the weighted averages (95% CI) of every plant inoculated with one of three example rhizobial strains (3B, Strain #2; 3C, Strain #14; 3D, Strain #24) in well-watered and drought treatments (N = 12 plants/strain/watering treatment). All plots are derived from the same RDA model.

Rhizobial strains induced unique multivariate plant trait phenotypes independent of environment (Fig. 4A: position of ellipses: F_1,23_ = 1.76, *p* = 0.001). Herbivory also induced differences in multivariate plant traits along the first RDA axis, as exposed plants were associated with increased photosynthesis and specific leaf area while non-exposed plants were more associated with increased photoprotection traits (Fig. 4A: black ellipses: F_1_ = 12.13, *p* = 0.001). Interestingly, strain-by-environment interactions drove variation in multivariate plant traits (Fig. 3B-D: position of ellipses: F_1,23_ = 1.23, *p* = 0.06) and trait plasticity under drought (Fig. 3B-D: size of ellipses: *betadisper*: d.f. = 47, F = 1.36, *p =* 0.07). For example, in strain #2, droughted plants shifted traits towards increased biomass along the second RDA axis and showed less plasticity in trait responses than well-watered plants (Fig. 4B). Well-watered plants inoculated with strain #14 were located centrally in the ordination space, but droughted plants shifted traits towards increased NPQt and leaf temperature differential; they also showed greater plasticity in their response to drought than well-watered plants (Fig. 4C). Plants inoculated with strain #24 were generally associated with increased root:shoot ratios in both watering treatments, but these plants exhibited reduced plasticity in their response to drought (Fig. 4D). Therefore, the effect of drought on shifts in multivariate plant trait expression and plasticity was also strain-dependent. A complete interaction figure displaying all twenty-four strains in each watering treatment is provided in the Supplementary Material (Fig. S2).

### How is plant performance and resource allocation impacted by rhizobial strain, drought, and herbivory?

The number of nodules per plant differed across rhizobial strains (Fig. 5; F_1,23_ = 9.01, *p* < 0.001). The removal of plants inoculated with strain #24 from the model did not change the significant strain effect on the number of nodules (F_1,22_ = 2.05, *p* = 0.003). However, we did not find evidence that rhizobial strain affected shoot or root plant biomass (Table S2). There was a 14% reduction in total plant biomass in ‘+ herbivory’ plants compared to ‘-herbivory’ plants (F_1,493_ = 22.7, *p* < 0.001). Weak evidence suggests that root:shoot ratio decreased with drought. This means that droughted plants allocated more biomass to their root systems, while well-watered plants allocated more biomass to their shoots (F_1,10_ = 3.75, *p* = 0.081). Additionally, there was no main or interactive effect of rhizobial strain identity, watering, or herbivory on chlorophyll content (Table S2).

**Figure 5.**
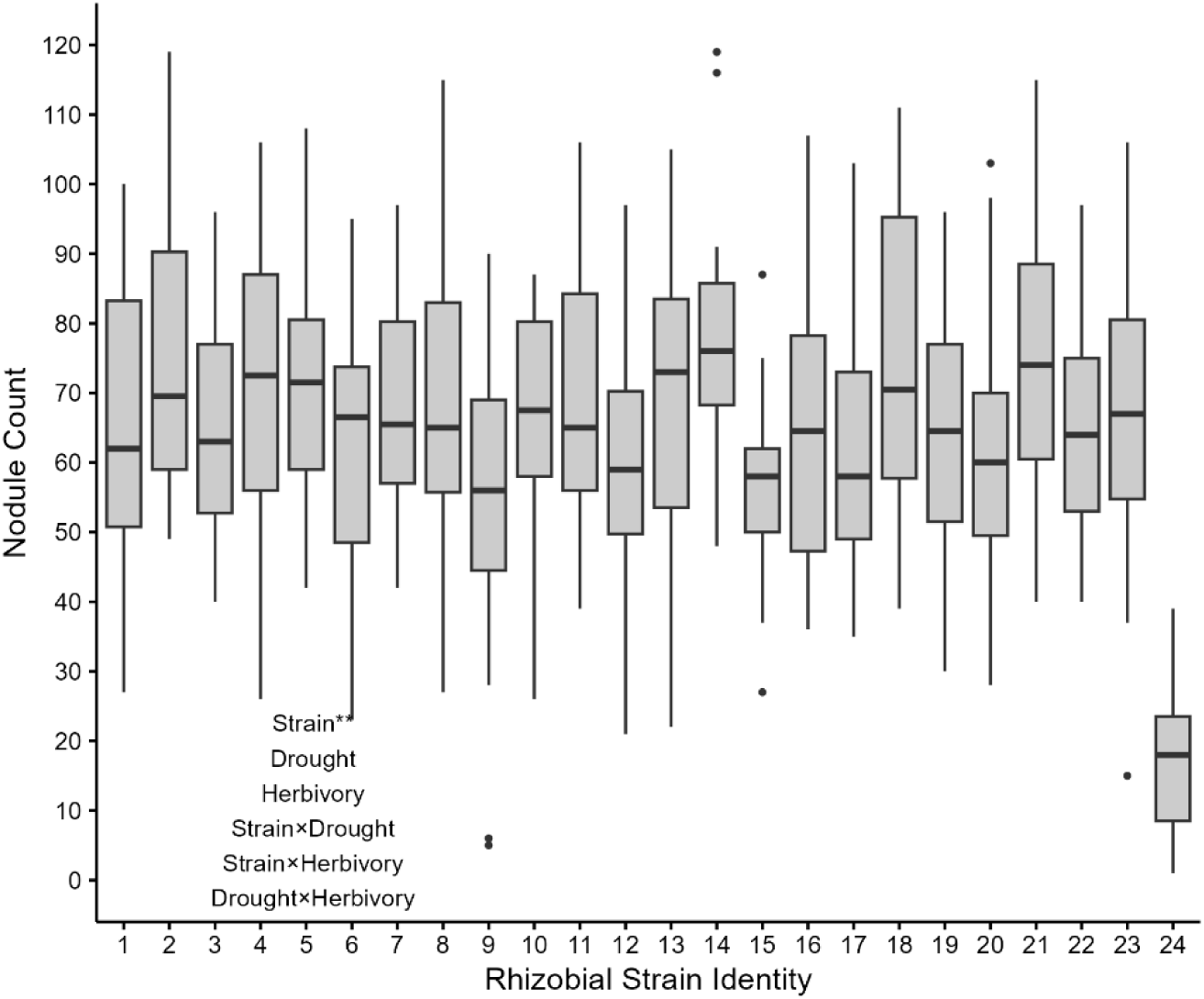
Rhizobial strains differed in the average number of nodules formed. Boxplot showing nodule count grouped by rhizobial strain (N = 24 plants/strain). ** = p ≤ 0.05.

## Discussion

We explored the role of genetically distinct rhizobial strains on plant functional traits and performance under multiple stressors, and how these effects cascaded up to higher trophic level interactions. We show that drought-induced changes in univariate and multivariate soybean functional trait syndromes are influenced by rhizobial partner genotype (G_Strain_×E_Drought_). Caterpillar feeding further modified photosynthetic and photoprotection traits both independently (E_Herbivory_) and in concert with drought (E_Drought_×E_Herbivory_). Additionally, we observed strain-specific responses to drought that indirectly affected caterpillar growth rates, but not leaf damage. We observed strain-dependent changes in plant nodulation and root:shoot ratio, but not chlorophyll content. Overall, we conclude that genetic variation among symbiotic partners interacts with abiotic and biotic stress to influence ecological interactions. Our work describes the context-dependency of tripartite species interactions, and how they are shaped by genetic, abiotic, and biotic factors.

We found that variation in caterpillar growth rates under drought was rhizobial strain-dependent (Fig. 2), but herbivory was not. Insect herbivores do not encounter a single plant trait when feeding. Instead, they experience a combination of plant traits, which may broadly influence herbivore growth rates (Laughlin & Messier, 2015; Burghardt, 2016; Wetzel et al., 2016). Herbivore growth rate integrates several herbivore traits, such as assimilation efficiency and palatability, making it a useful metric for understanding herbivore performance and plant quality (Scriber & Feeny, 1979; Coley et al., 2006). In our experiment, we found evidence of changes in resource allocation between strains and watering treatments in a multivariate analysis (Fig. 4), which could change palatability and tissue processing for herbivores. Together, these results indicate that differences in plant quality influenced caterpillar growth rate variation more than caterpillar intake rate.

Plants modify their functional traits to adapt to their environment (Grime et al., 2006; Reich, 2014; Heilmeier et al., 2019). Evidence from other legume systems demonstrates that rhizobia positively influence plant performance under drought and in well-watered environments (Staudinger et al., 2016; Bustos-Segura et al., 2024).

However, little work has addressed whether genetic variation of rhizobial partners influences plant functional traits that alter stress phenotypes under combined stress environments. We observed that strain identity mediated the effects of drought on plant photosynthetic and photoprotection traits (Fig. 3). Some strains enhanced photosynthesis or photoprotection under drought compared to others, indicating strain-based variation in plant drought tolerance. Shifting growth and tolerance traits under stress may also be explained by variance in N provisioning (Cerezini et al., 2020; Freitas et al., 2022). Future work is required to understand whether the effects of individual strain-based variation on plant trait plasticity and plant-insect interactions remain consistent under field conditions.

Different rhizobial strains induced distinct plant stress phenotypes (Fig. 4). A significant G×E interaction between rhizobial strain identities and drought on multivariate whole-plant trait expression, indicated by the difference in the position of ellipses (Fig. 4), suggests that strains differentially induce distinct trait syndromes under well-watered and drought conditions. Some strains induced shifts towards increased photosynthetic traits under drought, while others induced shifts towards plant protection. Furthermore, a significant G×E interaction between rhizobial strains and drought on multivariate whole-plant trait plasticity, reflected by the shapes of ellipses (Fig. 4B-D), indicates that strains either enhanced or constrained trait plasticity during stress. Our results highlight the role of microbial symbionts in shaping plant responses to stress, which may alter plant strategies and tradeoffs between protection and growth in stressful environments.

Our work expands on previous studies describing the role of rhizobia in influencing plant outcomes across variable stress environments by altering fixed N allocation and enhancing plant stress recovery (Staudinger et al., 2016; Thompson & Lamp, 2021; Bustos-Segura et al., 2024). Additionally, our work builds on previous studies that examined the role of rhizobial presence on higher trophic level interactions, particularly in reducing herbivore performance and herbivory (Ballhorn et al., 2007; Katayama et al., 2011; Thamer et al., 2011). Multiple strains can also establish and compete for resources from the same plant in many potential unique combinations (Jiao et al., 2015; La Pierre et al., 2017; Taylor et al., 2020; Batstone et al., 2023). Biotic interactors from higher trophic levels, such as herbivores, have been recently shown to positively or negatively shape diversity effects on symbiotic outcomes (Duffy et al., 2002; van der Plas et al., 2019). A recent study also points to the role of rhizobial partner diversity in reducing subsequent plant-insect interactions (Komatsu et al., 2023). Our research builds on this framework by showing that genetic variation between symbiotic microbial partners can not only differentially influence plant functional trait expression across environments, but that these effects can also impact higher order multitrophic interactions.

In our study, we emphasize the importance of rhizobial genetic variation in mediating multitrophic interactions in variable environments. We specifically wish to highlight that plant trait variation and the ability to induce changes in plant traits and, consequently, plant-insect interactions, can exist within a single rhizobial population (G), and the degree to which environment matters (G×E). In soils, there is a large amount of genetic variation of rhizobia, or strains, whose role in influencing plant plasticity to environmental stressors, multitrophic interactions, and ecosystem functioning has been scarcely examined. Future work should also examine whether there are differences in strain-generated changes in interaction outcomes across multiple plant genotypes (G_Strain_×G_Plant_).

Our results have key implications for the management of symbioses in agroecosystems. Drought during the growing season leads to more frequent pest outbreaks (Rosenweig et al., 2001; Cornelissen et al., 2011), which strongly reduces crop yields and quality (Frederick et al., 2001; Mohammadi et al., 2012; Zipper et al., 2016). Developing rhizobial inoculants for use in commercial farming operations often relies on inoculating field soils with a single strain, selected based upon plant yield in a single environmental context (Kaminsky et al., 2019). Our results indicate that these decisions should be more nuanced to account for tripartite species interactions under varying environmental conditions. For example, in our study, one strain did not lead to consistently high expression of functional traits and resistance to herbivory across combined stressors; rather, we found some strains influenced particular traits and herbivore performance more strongly than others, which often depended on the environmental context where the interaction took place. These results suggest that when developing potential solutions to agricultural sustainability under climate change, it is unlikely that one strain will be found that can be commercially reared and applied to provide high yields or stress-mitigating benefits under all environmental conditions.

Instead, our results highlight the need for incorporating and bolstering genotypically diverse, native rhizobial communities that communally protect leguminous crops under increasing environmental stress (Saad et al., 2020; Mendoza-Suárez et al., 2021; Poppeliers et al., 2023).

## Acknowledgements

We thank Jamie Pullen, Lauren Schmitt, and Elizabeth Butz for assisting with field and lab research. We thank Megan Holbert and Sydney Wallace for technical assistance at the UMD Research Greenhouse Complex. We thank the UMD Soybean Variety Trial Network for access to field plots. We thank Liana T. Burghardt for valuable comments and feedback on the manuscript. This project was supported by Agriculture and Food Research Initiative Competitive Grant no. 2019-67013-29139 from the US Department of Agriculture.

## Data Accessibility

All archived data is available from the FigShare repository: https://figshare.com/s/802d36026fe98d17976b

## Author Contributions

KJK, JDP, and KTB obtained funding for the research. BAR, KJK, JDP, KM, CTH, SJMA, and KTB conceived the ideas. BAR, KJK, JDP, KM, SJMA, and KTB designed methodology. BAR, KM, SJMA, and CTH collected the data. BAR analyzed the data and led the writing of the manuscript. All authors contributed critically to the drafts and gave final approval for publication.

