## Supplement file for "Microbial partners drive legume trait plasticity and tripartite interaction outcomes under combined stress environments"

Authors:
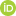
 Brendan A. Randall^1^, Kimberly J. Komatsu^2,3^, John D. Parker^3^, Kelsey McGurrin^1^, Sarah J. M. Alley^3^, Chase T. Hearn^1^, Karin T. Burghardt^1^

^1^Department of Entomology, University of Maryland, College Park, MD 20742 USA

^2^Department of Biology, University of North Carolina at Greensboro, Greensboro, NC 27402 USA

^3^Smithsonian Environmental Research Center, 647 Contees Wharf Road, Edgewater, MD 21037 USA

| **Trait** | **Description** |
| --- | --- |
| Phi2 | Maximum quantum yield of Photosystem II as measured by PhotosynQ according to Kulhgert et al., 2016. |
| PhiNO | Non-regulatory heat loss as measured by PhotosynQ according to Kuhlgert et al., 2016 |
| NPQt | Non photochemical quenching as measured by PhotosynQ according to Tietz et al., 2017; Kuhlgert et al., 2016 |
| LTD | Leaf temperature differential (ambient temp-leaf temp) |
| SLA | Specific leaf area (mm^2^/mg) according to Vile et al., 2005 |
| LWC | Relative leaf % water content according to Vile et al., 2005 |
| Chlorophyll | As measured by PhotosynQ according to Kuhlgert et al., 2016 |
| Height.cm | Plant height measured eight weeks after planting |
| Shoot.g | Shoot biomass harvested (g) |
| Root.g | Root biomass harvested (g) |
| RS.ratio | Proportion of total biomass in root tissues compared to shoot tissues |
| Nodule.count | Number of nodules harvested from each plant |

Table S1. Summary and descriptions of the twelve plant physiological and performance plant traits used in RDA.

| **Trait** | **Predictor** | **df** | ***F* value** | ***p*-value** |
| --- | --- | --- | --- | --- |
| Relative growth rate (mg/day-1) | Strain | 23 | 1.10 | 0.351 |
|  | Watering | 1 | 0.42 | 0.529 |
|  | Strain:Watering | 23 | 1.53 | 0.063* |
| Consumed leaf area (cm^2^) | Strain | 23 | 1.01 | 0.454 |
|  | Watering | 1 | 0.79 | 0.395 |
|  | Strain:Watering | 23 | 0.86 | 0.649 |
| Proportion leaf area removed (%) | Strain | 23 | 0.94 | 0.551 |
|  | Watering | 1 | 0.57 | 0.467 |
|  | Strain:Watering | 23 | 0.88 | 0.628 |
| Phi2 (*Δφ_F_/φF_m_*) | Strain | 23 | 1.67 | **0.033** |
|  | Watering | 1 | 0.17 | 0.700 |
|  | Herbivory | 1 | 4.93 | **0.026** |
|  | Strain:Watering | 23 | 2.30 | **< 0.001** |
|  | Watering:Herbivory | 1 | 1.50 | 0.201 |
| PhiNO (F*_S_*/F*_m_*) | Strain | 23 | 1.19 | 0.243 |
|  | Watering | 1 | 0.03 | 0.862 |
|  | Herbivory | 1 | 12.85 | **< 0.001** |
|  | Strain:Watering | 23 | 2.38 | **< 0.001** |
|  | Watering:Herbivory | 1 | 4.10 | **0.042** |
| NPQt | Strain | 23 | 1.56 | **0.049** |
|  | Watering | 1 | 0.03 | 0.858 |
|  | Herbivory | 1 | 7.47 | **0.007** |
|  | Strain:Watering | 23 | 2.20 | **0.001** |
|  | Watering:Herbivory | 1 | 2.07 | 0.151 |
| Leaf temperature differential (°C) | Strain | 23 | 1.52 | 0.058* |
|  | Watering | 1 | 0.11 | 0.746 |
|  | Herbivory | 1 | 2.82 | 0.094* |
|  | Strain:Watering | 23 | 0.66 | 0.886 |
|  | Watering:Herbivory | 1 | 1.90 | 0.169 |
| Specific leaf area (mm^2^/mg) | Strain | 23 | 0.99 | 0.473 |
|  | Watering | 1 | 0.55 | 0.475 |
|  | Herbivory | 1 | 7.95 | **0.005** |
|  | Strain:Watering | 23 | 0.93 | 0.555 |
|  | Watering:Herbivory | 1 | 0.31 | 0.580 |
| Relative leaf water content (%) | Strain | 23 | 1.18 | 0.261 |
|  | Watering | 1 | 0.10 | 0.760 |
|  | Herbivory | 1 | 11.5 | **< 0.001** |
|  | Strain:Watering | 23 | 1.24 | 0.202 |
|  | Watering:Herbivory | 1 | 0.02 | 0.876 |
| Relative chlorophyll (SPAD) | Strain | 23 | 0.96 | 0.531 |
|  | Watering | 1 | 0.22 | 0.654 |
|  | Herbivory | 1 | 2.09 | 0.132 |
|  | Strain:Watering | 23 | 0.71 | 0.818 |
|  | Watering:Herbivory | 1 | 2.46 | 0.102 |
| Plant height (cm) | Strain | 23 | 1.23 | 0.216 |
|  | Watering | 1 | 105.1 | **0.009** |
|  | Herbivory | 1 | 0.50 | 0.480 |
|  | Strain:Watering | 23 | 0.98 | 0.492 |
|  | Watering:Herbivory | 1 | 1.65 | 0.200 |
| Shoot biomass (g) | Strain | 23 | 1.33 | 0.141 |
|  | Watering | 1 | 1.24 | 0.291 |
|  | Herbivory | 1 | 52.6 | **< 0.001** |
|  | Strain:Watering | 23 | 0.68 | 0.867 |
|  | Watering:Herbivory | 1 | 0.75 | 0.251 |
| Root biomass (g) | Strain | 23 | 0.87 | 0.638 |
|  | Watering | 1 | 6.90 | **0.025** |
|  | Herbivory | 1 | 0.61 | 0.433 |
|  | Strain:Watering | 23 | 0.85 | 0.669 |
|  | Watering:Herbivory | 1 | 1.74 | 0.188 |
| Root:shoot ratio | Strain | 23 | 1.17 | 0.262 |
|  | Watering | 1 | 3.75 | 0.081* |
|  | Herbivory | 1 | 27.2 | **< 0.001** |
|  | Strain:Watering | 23 | 0.82 | 0.706 |
|  | Watering:Herbivory | 1 | 0.56 | 0.454 |
| Nodule count (#/plant) | Strain | 22 | 2.05 | **< 0.001** |
|  | Watering | 1 | 4.59 | 0.180 |
|  | Herbivory | 1 | 0.21 | 0.607 |
|  | Strain:Watering | 22 | 1.32 | 0.159 |
|  | Watering:Herbivory | 1 | 0.44 | 0.504 |

Table S2. Summary table of univariate linear mixed effects model results for three herbivore (caterpillar relative growth rate, consumed leaf area, and proportion of leaf area removed) and twelve plant physiological and performance traits. P-values are emboldened if *p* ≤ 0.05 and starred (*) if 0.05 < *p* < 0.1.

| Predictor | df | Variance | Pseudo-F | *p*-value |
| --- | --- | --- | --- | --- |
| Strain | 23 | 0.71 | 1.76 | **0.001** |
| Herbivory | 1 | 0.21 | 12.13 | **0.001** |
| Strain:Watering | 23 | 0.48 | 1.20 | 0.06* |
| Watering:Herbivory | 1 | 0.03 | 1.56 | 0.13 |

Table S3. Summary table of a simplified distance-based redundancy analysis (db-RDA) model results for twelve plant physiological and performance traits (Phi2, PhiNO, NPQt, leaf temperature differential, SLA, LWC, relative chlorophyll, plant height, shoot and root biomass, root:shoot ratio, and nodule count) by experimental predictors. Main effect of watering was aliased from the model because it was highly correlated with our conditioning “block” term. The strain:herbivory interaction was removed from the model because it explained virtually no variance. The simplified model did not change any of the model results. P-values are emboldened if *p* ≤ 0.05 and starred (*) if 0.05 < *p* < 0.1.

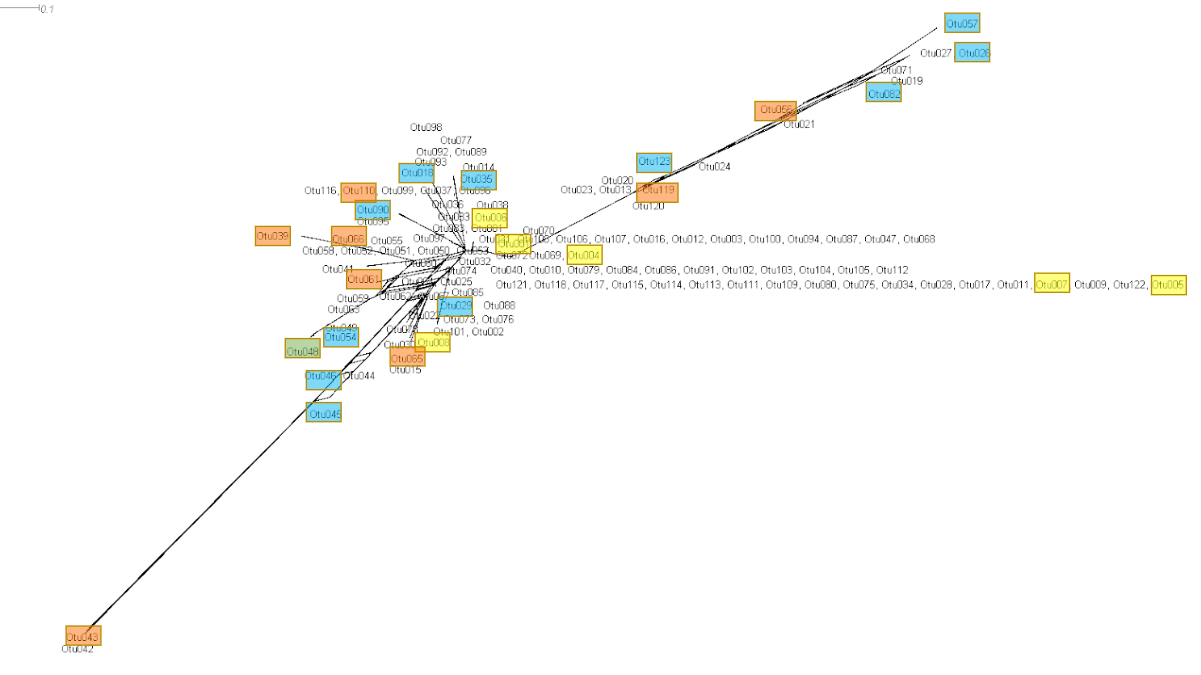

Figure S1. Neighborhood network diagram of 123 rhizobia OTUs collected and sequenced from a rhizobial trapping greenhouse experiment. OTUs that were centrally clustered in space were more closely related to each other, while OTUs on separate branches were more distantly related to each other. Twenty-four OTUs were selected for the single-strain manipulation experiment based on several criteria, with the aim of capturing a representative sample of genetic differences and environments: OTUs from distinct branches, OTUs that were dominant and were collected multiple times (yellow boxes), as well as OTUs collected from soybeans grown in ambient (blue boxes) and drought (orange boxes) environments. The selected OTUs were classified within the *Bradyrhizobium diazoefficiens* (N=21) and *Bradyrhizobium elkanii* (N=3) species complexes.

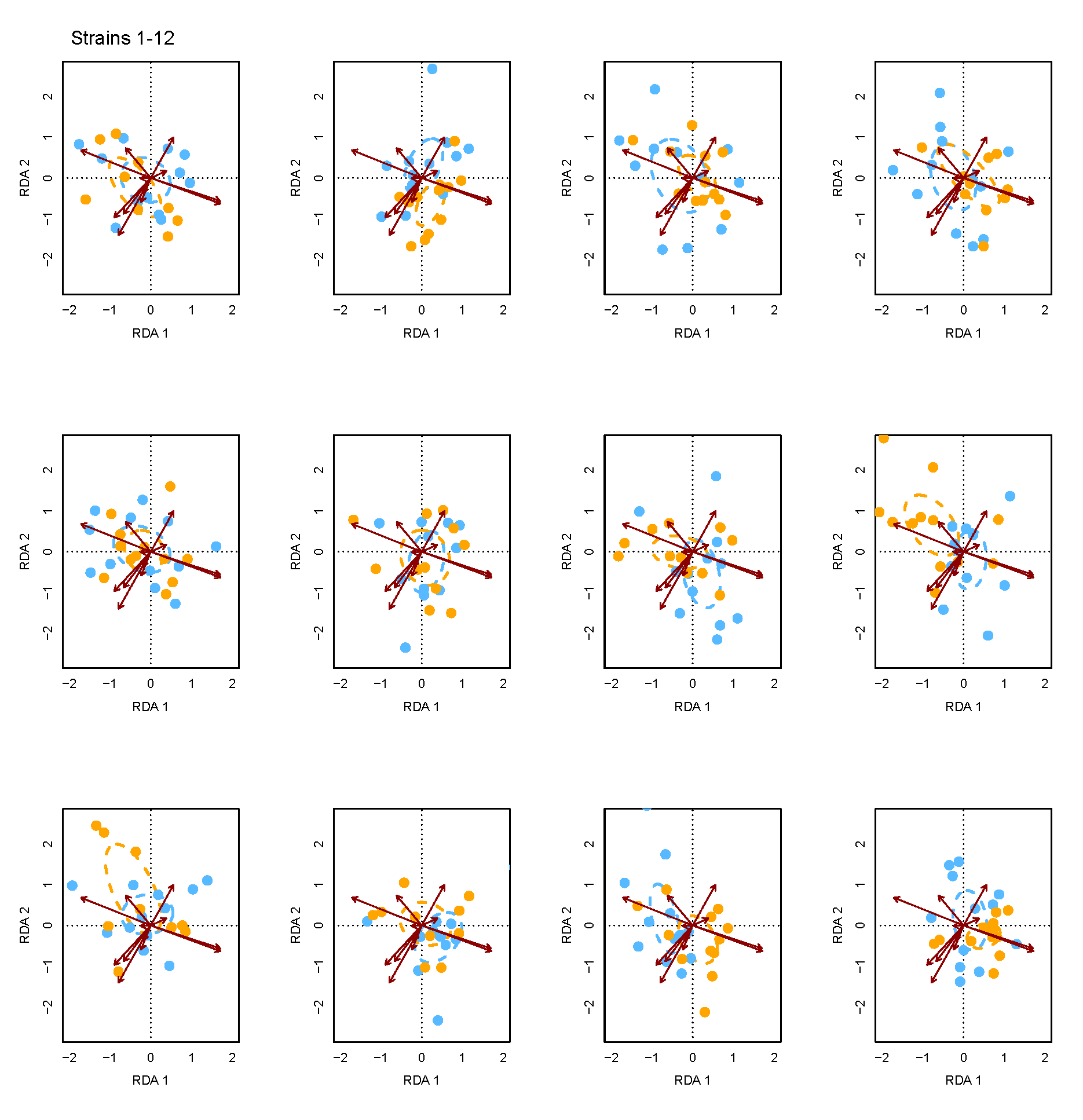

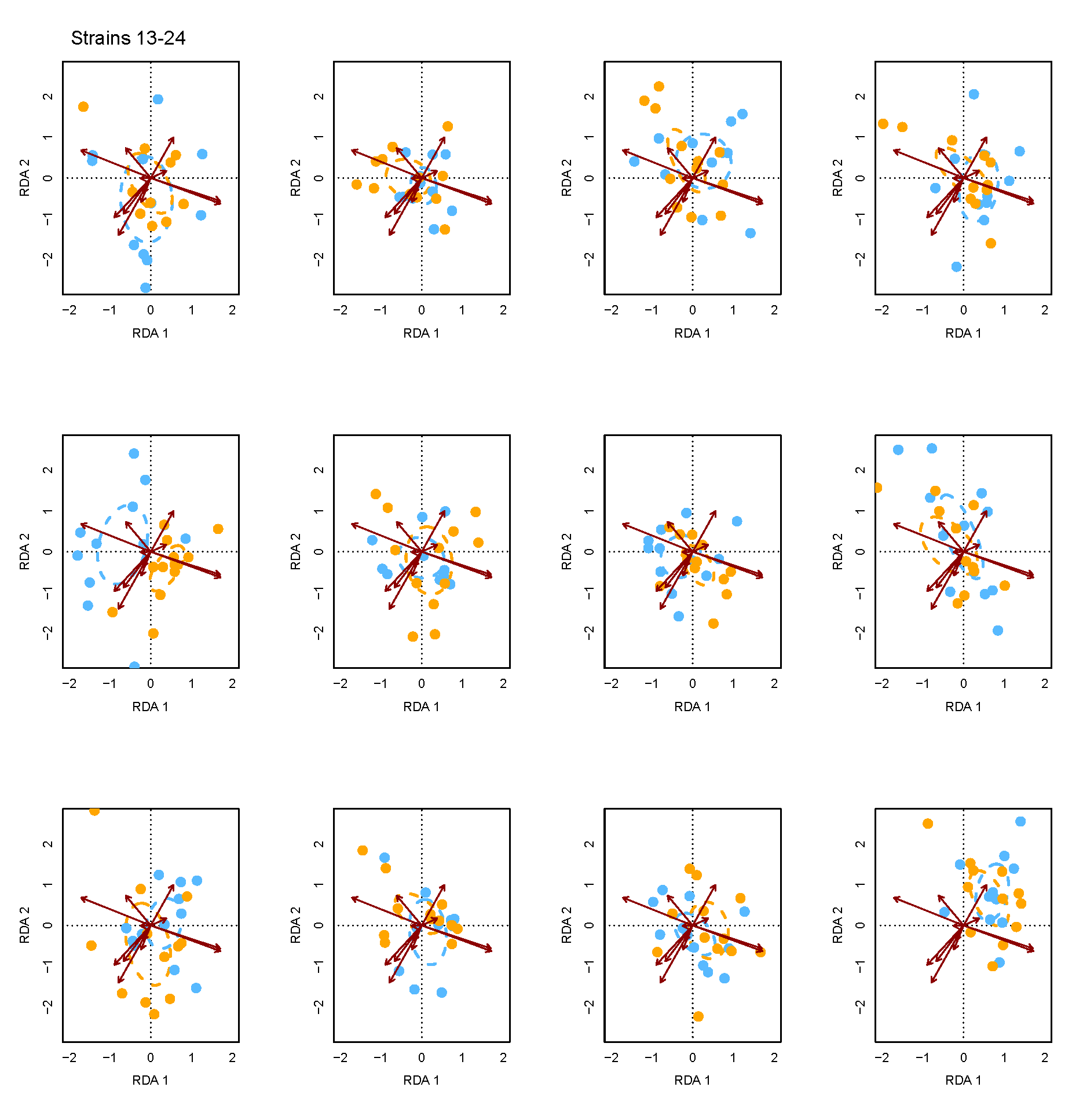

Figure S2. RDA plot with all twenty-four strains included, with plants grouped by the strain they were inoculated with (N=24). Colored ellipses represent the weighted averages (95% CI) of all plants inoculated with a particular strain that were exposed to well-watered (blue ellipses) or drought (orange ellipses) environments. Individual plants for each strain identity and watering treatment are represented as dots.
